# Dissipation During the Gating Cycle of the Bacterial Mechanosensitive Ion Channel Approaches the Landauer’s Limit

**DOI:** 10.1101/2020.06.26.174649

**Authors:** Uǧur Çetiner, Oren Raz, Sergei Sukharev

## Abstract

The Landauer’s principle sets a thermodynamic bound of *k_B_T* ln 2 on the energetic cost of erasing each bit of information. It holds for any memory device, regardless of its physical implementation. It was recently shown that carefully built artificial devices can saturate this bound. In contrast, biological computation-like processes, e.g., DNA replication, transcription and translation use an order of magnitude more than their Landauer’s minimum. Here we show that saturating the Landauer bound is nevertheless possible with biological devices. This is done using a mechanosensitive channel of small conductance (MscS) from *E. coli* as a memory bit. MscS is a fast-acting osmolyte release valve adjusting turgor pressure inside the cell. Our patch-clamp experiments and data analysis demonstrate that under a slow switching regime, the heat dissipation in the course of tension-driven gating transitions in MscS closely approaches its Landauer’s limit. We discuss the biological implications of this physical trait.

## I. INTRODUCTION

Any computation done on a physical system is subject to fundamental limitations imposed by the laws of physics. For example, the uncertainty principle implies that in order to perform an elementary logical operation faster than some Δ*t*, at least an average amount of energy *E* ≥ *πħ*/2Δ*t* must be consumed^1^. Another important bound on computation, which is at the main focus of this work, is set by the laws of thermodynamics. According to the Landauer’s principle^2^, at least *k_B_T* ln 2 of heat dissipation must accompany any one-bit erasing process.

Here *k_B_* is the Boltzmann constant and *T* is the ambient temperature. Equality can be achieved for quasi-static (reversible) erasure protocols. The heat released into the environment during the erasure of information assures that the total increase in the entropy of the system and bath together is a non-negative quantity. Importantly, this bound applies to any non-reversible erasing process of a memory, regardless of the physical system that was used to implement it. Therefore, the Landauer’s principle demonstrates the interplay between physics and information. In recent decades, the Landauer’s principle was generalized to include: a probabilistic erasure process^3,4^; other types of thermodynamic resources^5^; entropically unbalanced bits^6^; a unified view on the cost of erasing and measuring a bit^7,8^; *N* state bit^9^; optimal erasure at finite time^10,11^ and others^12^.

The existence of a fundamental bound does not imply that the bound can be saturated. And indeed, current computer memory devices dissipate about six orders of magnitude more energy than the minimal amount required by the bound. Similarly, estimations of the energy dissipated in biological computations such as DNA replication, transcription and translation show that these are done with one to two orders of magnitude more dissipation than required by the Landauer’s bound^13^. Recently, however, it was demonstrated that carefully built artificial systems can actually operate very close to the Landauer’s bound. This was achieved with several types of systems: a single colloidal particle in an optical^14,15^ or feedback^16–18^ traps, nanomagnetic bits^19–21^, superconducting flux bit^22^ and even quantum systems^23,24^. Based on these results, it is natural to ask whether there are any biological memory erasing processes that operate close to the Landauer’s limit.

Out of many biological systems, the bacterial mechanosensitive ion channels of small and large conductance, MscS and MscL, appear to be the most tractable two-state (closed open) systems controlled by tension in the surrounding membrane^25–27^. They function as osmolyte release valves when bacteria face changing environmental osmotic conditions, such as in the rain. While the large-conductance MscL channel opens by extreme near-lytic tensions and acts as an emergency valve, the small-conductance MscS channel opens at moderate tensions and appears to be active throughout the normal bacterial lifecycle^26,28–30^.

In this work, we present a framework for analysis of heat dissipation in membrane channels gated by tension. We employ the patch-clamp technique applied to the native *E. coli* membrane to record discrete single-molecule opening and closing events in MscS under specially designed tension stimuli and extract the dissipated heat that accompanies gating transitions. The state of the ion channel, which can be either “open” or “closed”, encodes a single bit of information. Setting the experimental conditions such that the channel occupies these two states with equal probability introduces the maximum degree of randomness. Changing the biasing tension that re-distributes the channel population to one particular state is equivalent to “erasing the memory” stored in the initially randomized population. We extract the heat dissipated during the “restore to open” process imposed with different rates and show that this system dissipates substantially at high transition rates, but under slower driving protocols, MscS gating closely approaches its Landauer’s limit. We discuss the physiological importance of this physical trait, which predicts the activation of MscS with minimal dissipation under moderate osmotic shocks experienced by bacteria.

## II. EXPERIMENTAL AND THEORETICAL SETUP

To measure the dissipated heat during the erasure of a single bit, Landauer suggested to use a “restore to one” protocol^2^, which results in the bit occupying a single state – the “one” state, regardless of the initial state of the bit. He then argued that the heat dissipated in applying this protocol, averaged over the two initial states of the bit, must be at least *k_B_T* ln 2.

Our experiments use the patch-clamp technique^31–33^ to implement the “restore to one” protocol on MscS ion channels, where the one-bit information is stored in the “open” and “closed” states of a single MscS ion channel. To this end, a piece of *E. coli*’s inner membrane harboring about 20 MscS channels is sealed to the polished tip of a glass micropipette. Because the native inner membrane of wild-type *E. coli* contains a mixed population of hundreds of MscS-type and MscL channels, we used a tightly-controlled expression system (see materials and methods) in a strain devoid of seven endogenous mechanosensitive channel genes (Δ *mscL, mscS, mscK, ybdG, ynal, ybiO,* and *yjeP*). Moderate induction produced on average 20 channels per patch, which was optimal for the recording and analysis. The micropipette with the clamped membrane is immersed into the bath solution at room temperature. The high resistance of the glass-membrane seal provides an electrical isolation of the patch separating two aqueous solutions (Fig.1 left). Application of suction to the pipette generates tension in the patch and activates the channels (see SI for the details of tension calibration). These molecular activation events can be monitored with pico-ampere precision, which allows us to track the distribution of channels between the open and closed states at a single-channel resolution^34^ (see, for example, Fig. 2B). In a typical experiment, after seal formation and patch excision, a linear ramp of negative pressure (suction) from zero to the saturating level is applied to the patch with simultaneous recording of the population current. This step determines the activation pressure midpoint (*p*_0.5_) at which the population is equally distributed between the closed and open conformations, i.e. the state of highest uncertainty. In the following “bit erasure” protocol, the pressure is quickly ramped to *p*_0.5_, the population is allowed to equilibrate for 3 s and then the pressure is ramped with different rates to a higher level, where all channels uniformly assume the open conformation (state of highest certainty). The recorded traces with easily discernable single-channel steps are analyzed with the “edge detection” protocol^35^ as described below.

**FIG. 1:**
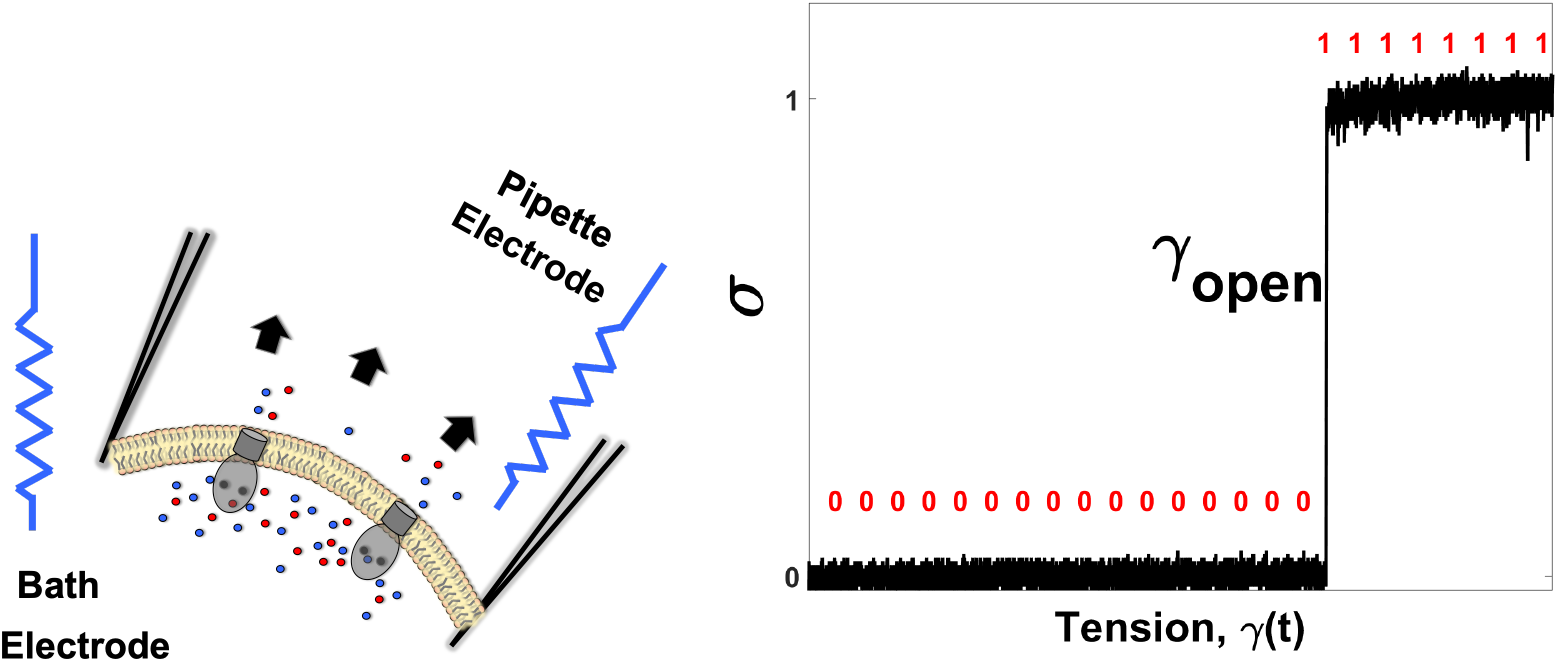
Left: Schematic description of the experimental apparatus. The Giga-ohm resistance of the piece of *E. coli*’s inner membrane with naturally embedded mechanosensitive ion channels (MscS) seals the micro-pipette, and provides an electrical isolation between its inner and outer sides. Application of suction to the glass pipette stretches the curved membrane according to Laplace’s law. This tension can change the state of the MscS channels, generating a detectable conducting pathways between the two electrodes. Right: The state of the channel (*σ*) as a function of the membrane tension (*γ*). A single channel event is shown. The state of the channel can be monitored with a high temporal resolution. The transition from the closed (0) state to the open state (1) occurs at *γ*_open_.

**FIG. 2:**
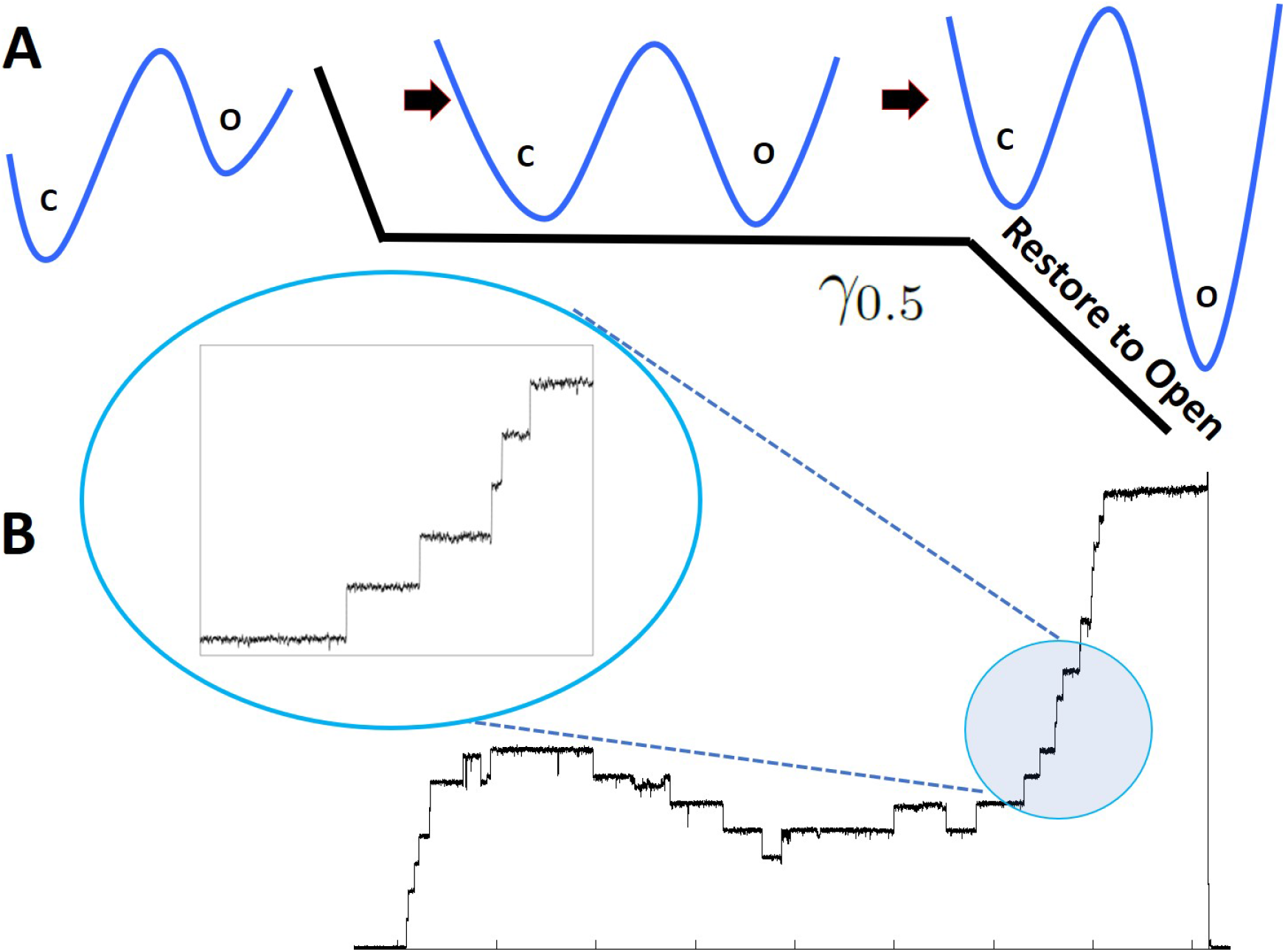
**A**: Restore to open protocol. When unperturbed, the channels naturally occupy the low energy configuration which is the closed state. In the first part of the protocol, the tension was quickly (0.25 *s*) increased to the midpoint tension *γ*_0.5_ (1.9 *k_B_T/nm*^2^)^42^ at which probability of finding a channel in the open or closed state is 0.5. The tension was kept fixed at *γ*_0.5_ for 3s to let the channels thermalize at this specific tension value. In the final setup, the tension was increased from *γ*_0.5_ to *γ_τ_* = *γ_max_* (3 *k_B_T/nm*^2^) in 0.25, 1, 5 and 10 seconds. Regardless of the initial state of the channels, at the end of the final step, all channels are forced to be in the open state. **B**: An experimental trace obtained from the restore to open protocol. In the final step, the tension was increased from *γ*_0.5_ to *γ_τ_* in 1s. The inset shows the single-channel gating events at a higher magnification during the restore to one operation.

At room temperature, the minimal dissipated heat set by the Landauer’s bound, *k_B_T* ln 2, is extremely low—about 10^−21^*joules*. This makes any direct measurement of the heat absorbed by the environment, e.g., by measuring its thermal expansion or temperature raising, highly challenging. Fortunately, the recent theory of stochastic thermodynamics^36^ suggests a way to measure the dissipated heat by watching the behavior of the thermal system itself, rather than measuring the environment. This method was used in measuring the saturation of the Landauer’s bound in artificial systems^14–20^. Principles of non-equilibrium statistical thermodynamics have been successfully applied to MscS ion channels in a different context^37^.

To discuss heat dissipation in the MscS ion channel, we model it as a two-state system (”open” and “closed”), and introduce a state variable, *σ* = 0 for a closed channel and *σ* = 1 for an open one. For a system with *N* such channels, we denote by 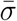 the average over the states of the di erent channels. In contrast to *σ* that can take only 0 and 1 as its value, 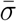 can take any value between 0 and 1 with an accuracy *N* ^−1^. Let *c*_closed_ and *c*_open_ denote the energies of the closed and open states of the ion channel itself. The total energy of the ion channels and the membrane is a function of the tension *γ* and the state variable *σ*, and is given by^37–39^:

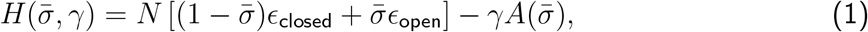

The additional term 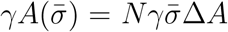 presents the decrease in the energy of the membrane that occurs when the channels open in response to the applied tension, *γ*. Here Δ*A* is the area difference between the closed and open state of the channel. Thus in the presence of external tension on the membrane, states with larger area become favorable^27,40^. The energy and area difference between the closed and open states of a single MscS channel were already measured in previous publications^37,40^, and are given by: Δϵ ☰ ϵ_open_ − ϵ_closed_ = 22 *k_B_T* and Δ*A* = 12 *nm*^2^. In what follows we used the standard assumption that Δ*A* and Δϵ are the same for all MscS channels.

Based on the above energy in the system, the total change in energy can be expressed as follows^41^:

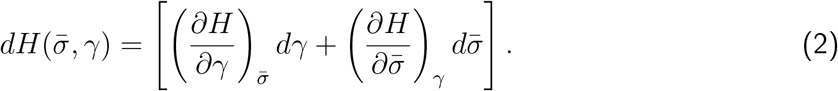

The first term on the square brackets of Eq.2 is the change in energy resulting from some external force changing the tension, which is therefore associated with work. The second term is the variation of the energy resulting from the change in the internal configuration of the system, namely due to redistribution between states. To conserve the total energy, this energetic change requires an exchange of energy with the surrounding thermal bath, and is therefore associated with heat. With these interpretations, the total heat and work associated with a realization of the experimental protocol can be written as:

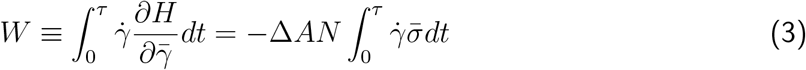

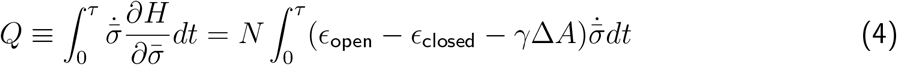

where *τ* is the protocol’s duration. In our experiments the tension *γ* is changed linearly with time, therefore we can write the work integral in the following form:

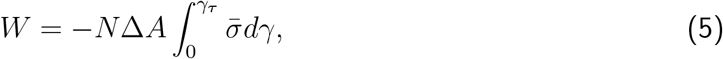

which can be interpreted as *N*Δ*A* times minus the area under the 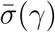 graph. The above definitions of heat and work imply the *N* → ∞ limit, to make sense of 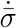. The tools of stochastic thermodynamics enable us to extend these definitions to small systems with even a single channel, where *σ* can take only discrete values of 0 or 1, and changes abruptly between them. In this case, the work in Eq.(5) can be directly used. To calculate the heat, however, we note that in this case *σ*(*t*), shown in Fig. 1, can be approximated with the Heaviside step function:

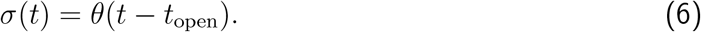

Exploiting the relations between the Heaviside step function and the Dirac delta function, we can write the heat integral in a particular realization:

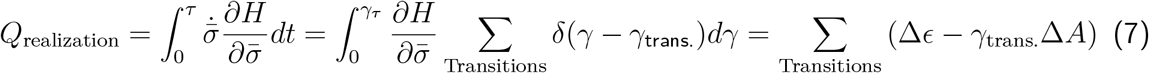

Alternatively, the heat can be expressed as the difference between the total change in energy and the work.

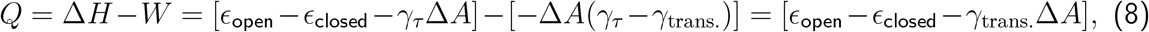

where we expressed 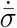 as a sum of delta functions located at the transition tensions *γ* trans. in this specific realization, and Δϵ and Δ *A* represent the changes corresponding to the specific transition, which can be both opening or closing of a channel. As the channels are independent, this definition also gives a heat value for each gating event. Stochastic thermodynamics assures^36^ that the average of the heat calculated by Eq.7 over many realizations converges to the correct ensemble average of the heat dissipation. Note that Eq. 7 with a minus sign corresponds to the heat released by the system into the environment: the difference between the intrinsic transition energy in the channel molecule (which is constant) and the work that is done on the molecule by external tension during the transition (which is proportional to applied tension) gives us the dissipated heat. By construction, the above definitions of heat and work recover both the first and second laws of thermodynamics^37^. With the above interpretation, heat and work can be associated with every realization of the protocol. It is important to note, however, that they do not have the same value at each single realization, and they may fluctuate from one realization to the next. Therefore, averaging over many trajectories is required to get a reliable estimate for the dissipated heat.

## III. RESULTS

In a typical experimental setup, the MscS channels naturally reside in the closed state when no pressure is applied on the system. Therefore, in the first stage of the experiment, we increased the membrane tension by applying suction pressure on the micro-pipette, to the midpoint tension value *γ*_0.5_ at which the probabilities of finding the channel in the closed or open states are equal, *P*_Open_ = *P*_Closed_ = 0.5 (this ramping is done during 0.25 *sec.*). We then let the system thermalize at this tension value (*γ*_0.5_) by keeping the pressure fixed for 3 *sec.* The system’s entropy per channel at this stage can be calculated as: *S*_Initial_ = −*k_B_* ∑_*i*_*P_i_* ln *P_i_* = *k_B_* ln 2 (See Fig. 2A)

In the second stage of the experiment, we increased the membrane tension to 3 *k_B_T/nm*^2^ at which *P*_Open_ ∼ 1. This was done at various ramping rates. This protocol mimics Landauer’s “restore to one” operation, which deletes a single bit of information. To see why, note that the channels are *restored* to the open state from an initial configuration where the close and open states are equally likely to be occupied. Since the channels are forced to the open configuration regardless of their initial status, this protocol is equivalent to the “restore to open”. The entropy of the system after this stage is given by: *S*_Final_ = −*k_B_* ∑_*i*_*P_i_* ln *P_i_* = 0. Therefore, the change in the entropy of the system is Δ*S* = *S*_Final_ − *S*_Initial_ = −*k_B_* ln 2. This operation corresponds to deleting a single bit of information^43^. To compensate for the system’s entropy decrease, the heat released into the environment must be at least *k_B_T* ln 2, otherwise the total entropy of the system and the bath decreases, leading to a violation of the second law of thermodynamic.

We repeated the above experiments many times and gathered ~200 single-channel events for each erasure protocol. In each realization, we monitored the heat released into the environment using Eq. (7) and the known values of Δϵ and Δ*A*. These were plotted as a function of the rate at which the tension was changed from *γ*_0.5_ to *γ_τ_* in Fig.(3). As expected, the averaged dissipated heat decreases with the protocol duration. At the slowest experimental erasure protocol achievable (see discussion), we reach very close to the Landauer limit of *k_B_T* ln 2, much closer than any other biological system reported so far.

**FIG. 3:**
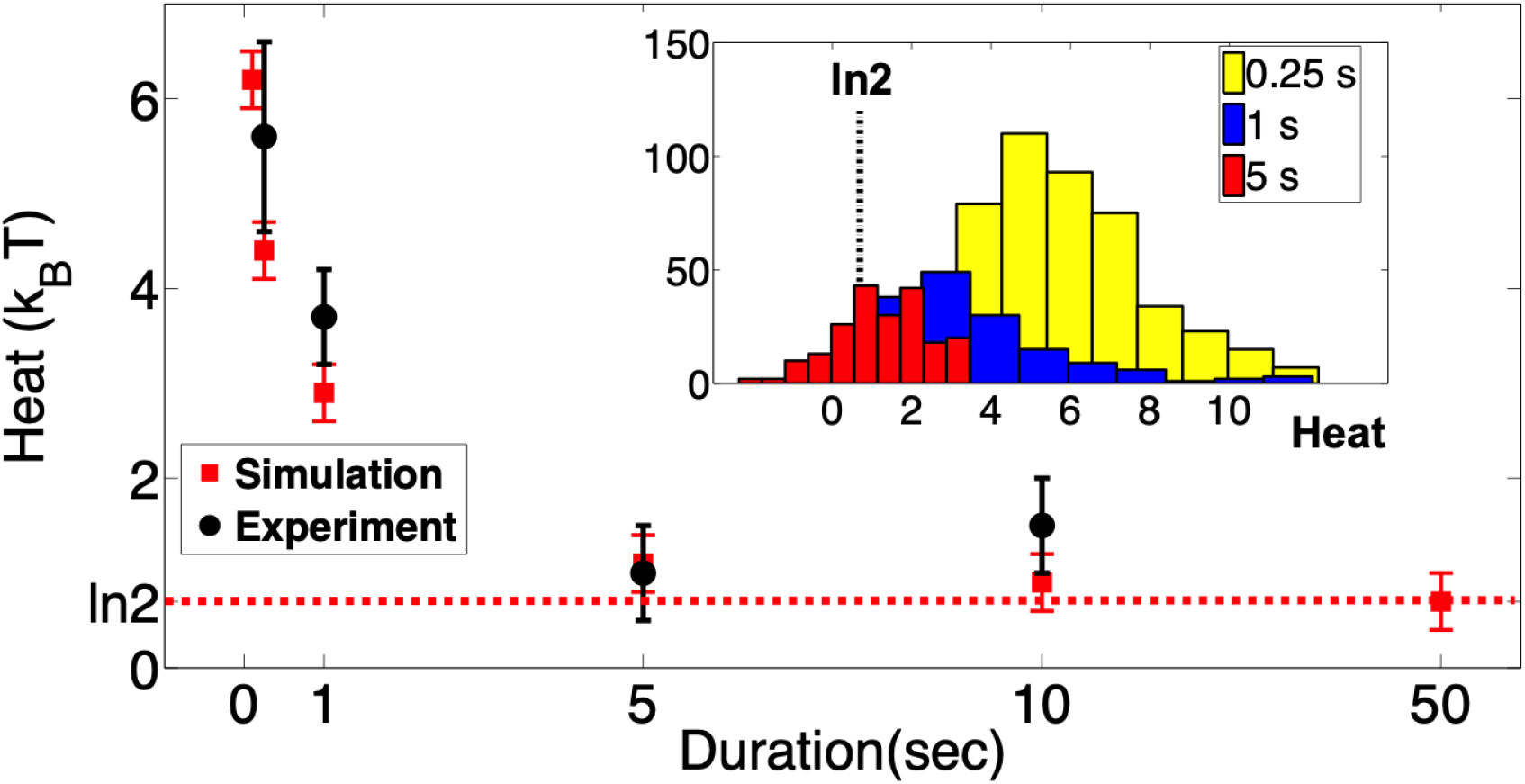
The average dissipated heat as function of the “restore to open” operation rate. As the channels are restored to the open state slower and slower (the duration increases), the average heat dissipated decreases, but it is always above the Landauer’s limit of *ln*2. Under sustained mechanical stimuli, the MscS channels inactivate wherein they enter a non-conductive and tension-insensitive state. Therefore, the slowest experimentally achievable erasure duration was limited to 10s after which the channels display significant inactivation. A Markov model of two-state MscS has been also simulated using QUBexpress software with different rates of restore to open protocol (red data points). The simulation result not only agree with the experimental counterpart but also saturates at the same limit of *ln*2. The simulation parameters are provided in the SI. The inset shows the histograms of heat distributions from which the averages are obtained.

To further verify our results, we simulated a Markovian model of MscS gating using our experimental protocol (*restore to open*) as the input driving force in the simulation using QUBexpress software^44^. The parameters used in the simulation and details of two-state Markov model of MscS are given in the supplement. Since in the simulations erasure protocols can be made arbitrarily slow, we obtained the heat distribution as a function of longer erasure protocols (Fig. 2, red data points). The simulation results are in a good agreement with the experimental measurements at short protocols, and for longer protocols they in fact saturate the *k_B_T* ln 2 bound.

## IV. DISCUSSION

Living systems are inherently dissipative, especially as they execute multiple steps of chemical energy conversion, pump metabolites, produce mechanical work, or maintain constant temperature. The question that the researchers studying *structural information content and cellular computation* try to address is not about the total energy balance and dissipation, but rather about the energy consumption by the “cellular switchboard” itself that turns the cellular processes on and off, replicating information and thus making decisions. Previous analysis based on the generalized Landauer bound^13^ have suggested that protein synthesis, which is an RNA-guided non-random polymerization of amino acids, takes about an order of magnitude more energy than the amount of information stored in the sequence requires. Synthesis of DNA on a DNA template, according to estimations^13^, consumes about two orders of magnitude more energy than the Landauer bound predicts. The problem with these systems is that the energy provided by the splitting of deoxyribonucleotide triphosphates (dNTPs) is strictly coupled with each polymer extension step, which makes this chemical energy component inseparable from the purely entropic change of information content.

In this work, we studied the mechanosensitive ion channel of small conductance (MscS) from *E.coli* acting as tension-operated membrane valve requiring no chemical energy input. MscS evolved to release excess osmolyte from cells in response to osmotic water influx that causes the cell envelope to swell and stretch. Opening of the entire MS channel population during strong shock massively dissipates internal ions and osmolytes that can amount up to 5-10% of cellular dry weight^26^. This undoubtedly inflicts a substantial energy and metabolite loss by the cell that is trying to evade lysis at any cost. However, as our results show, the operation of MscS itself in a slow (nearly equilibrium) regime costs that minimum, exactly as the Landauer limit predicts.

Our experimental conditions allowed us to treat MscS as a two-state a memory device. By applying tension to the patch membrane, we forced the population of channels to change its state occupancy, from which we measured the thermodynamic cost of deleting a single bit of information. The heat dissipated during the bit-erasing transition to the singular open state was measured as the average difference between the intrinsic transition energy in the channel molecule and the work that is done on the molecule by external tension. The dissipated heat measured with a short “restore to open” time (0.25 s) exceeded 5 *k_B_T*, whereas at slow ramps it approached ln 2 *k_B_T*, corresponding to the Landauer’s bound.

The practical requirements we had to satisfy in our experiments were as follows. (i) Because MscS channels tend to inactivate when exposed to moderate tension (*γ*_0.5_) for a prolonged period of time, the time for the state restoration protocol cannot be arbitrarily slow. In order to stay in the two-state regime, we used a short (0.5 s) pressure ramp to *γ*_0.5_, a 3-s equilibration, and a variable duration “erasure ramp” that was limited to 10 s. The non-inactivating mechanosensitive ion channel of large conductance MscL, would also be a good system for dissipation analysis, but it gates at near-lytic tensions where membrane patches become unstable^26^. For this reason, MscL was not used. (ii) MscS expression level had to be carefully adjusted through the use of a tight-promoter expression system such that the number of channels per patch (10-20) was suitable for the edge detection analysis of individual transitions. With all these precautions, a small degree of adaptation and inactivation were still observed (expected to be around 10% for a 3 sec holding time at *γ*_0.5_), which gave rise to a non-monotonic current response shown in Fig. 2.

Why is the energetic cost of erasing the bit of information encoded in the ion channel as low as the theoretic bound? Although we can only speculate, from a biological point of view this trait seems natural. Under hyperosmotic conditions, bacteria accumulate ions and organic osmolytes to maintain a positive turgor pressure inside the cytoplasm. Moreover, bacteria maintain relatively high voltage across the cytoplasmic membrane (150-200 mV) as a part of electrochemical potential driving ATP synthesis^45,46^. The thermodynamic and kinetic stability of the closed state, therefore, are critical because thermally-driven random opening events would produce deleterious leakage and uncoupling of bacterial energetics. Thus, evolution has perfected the energy gap (~22 *k_B_T*) between the end states and the height of the separating barrier such that thermal energy does not produce spurious openings at rest during the lifespan of bacteria. However, in the event of a sudden osmotic downshock such as during a rainstorm, cellular osmolytes are quickly released through mechanosensitive ion channels in order to reduce the turgor pressure. The low-threshold MscS and the high-threshold MscL channels are responsible for the bulk of osmolyte exchange in *E.coli*, but each channel is specialized in handling different magnitudes of osmotic shocks. The 3-nS MscL is an emergency valve that opens abruptly at near-lytic tensions (~3.5 *k_B_T /nm*^2^) and jettisons the osmolytes non-selectively. The 1-nS MscS, on the other hand, operates at moderate tensions (~2 *k_B_T /nm*^2^), and effectively counteracts small osmotic shocks. These channels evolved to defend bacteria under different osmotic conditions, e.g., emergency vs non-emergency situations, and they perform more efficiently under certain timescales^40^.

Stopped-flow experiments revealed that the characteristic time scales of bacterial swelling in response to an abrupt dilution vary from seconds at low shocks (100-300 mOsm downshifts) to 100 milliseconds at stronger (600-1000 mOsm) shocks^26^. Such strong osmotic down-shock experiments yield the typical timescales at which an emergency valve operates in nature. MscS populations residing in the cytoplasmic membrane of a bacterium are usually able to meet the kinetic requirement, i.e., opening and helping to reduce the internal turgor pressure by quickly releasing the excessive osmotic gradient before water influx rips open the cell. However, MscS, not being a true emergency valve, is somewhat inefficient when it is forced to open under timescales of 30-50 ms that correspond to super-threshold tensions in the cytoplasmic membrane generated by fast dilution *in-vivo*. This dissipation at higher tensions (and rates) is a “tax” imposed by a relatively high transition barrier providing a “safety curb” that precludes spurious openings at low tensions. However, under moderate osmotic shock conditions, when tension buildup in the cytoplasmic membrane occurs within a time span of a few seconds, MscS performs a smooth action in non-dissipative manner, which is consistent with the in vivo role of MscS in the overall osmotic fitness of *E. coli*.

## ACKNOWLEDGMENTS

This research was performed while UC was a PhD student in the Maryland Biophysics program. UC was supported by the U.S. Department of Education GAANN Mathematics in Biology Fellowship. OR is the incumbent of the Shlomo and Michla Tomarin career development chair, and is supported by the Abramson Family Center for Young Scientists and by the Israel Science Foundation, Grant No. 950/19. The work was supported by NIH R21AI105655 and GM107652 grants to SS. UC thanks Ms. Stephanie Sansbury for cloning MscS into tightly-regulated pBAD for expression system.

## SUPPLEMENTARY INFORMATION

### Material and Methods

#### Preparation of giant spheroplasts and patch clamp

The giant spheroplasts of E. coli were prepared following the protocol described in^34^. 3 *ml* of the colony-derived culture was transferred into 27 *ml* of LB containing 0.06 *mg/ml* cephalexin, which selectively blocks septation. After 1.5-2 hours of shaking in the presence of cephalexin, 100 250 *μm* long filaments formed. Toward the end of the filamentous growth stage, induction with 0.001 % L-Arabinose was done for 0-20 mins which gave 1-15 channels per patch. The filaments were transferred into a hypertonic buffer containing 1 *M* sucrose and subjected to digestion by lysozyme (0.2 *mg/ml*) in the presence of 5 *mM* EDTA. As a result, filaments collapsed into spheres of 3 – 7 *μm* in diameter in 7-10 mins. The reaction was terminated by adding 20 *mM Mg*^2+^. Spheroplasts were separated from the rest of the reaction mixture by sedimentation through a one-step sucrose gradient. Borosilicate glass (Drummond 2-000-100) pipets 1 – 1.3 *μm* in diameter were used to form tight seals with the inner membrane. The MS channel activities were recorded via inside-out excised patch clamp method after expressing them in MJF641. The pipette solution had 200 *mM KCI*, 50 *mM MgCI*_2_, 5 *mM CaCI*_2_, 5 *mM HEPES*. The bath solution was the same as the pipette solution with 400 mM sucrose added. Both pipette and bath solution had the pH of 7.4. Traces were recorded using Clampex 10.3 software (MDS Analytical Technologies). Mechanical stimuli were delivered using a high-speed pressure clamp apparatus (HSPC-1; ALA Scientific Instruments).

#### Tension Calibration

The pressure (P) was converted to the tension (*γ*) using the following relation: *γ* = (*P/P*_0.5_)*γ*_0.5_ assuming the radius of curvature of the patch does not change in the range of pressures where the channels were active (P >40 mmHg) and the constant of proportionality between tension and pressure is *γ*_0.5_*/P*_0.5_^25,27,42^. The midpoint tension, *γ*_0.5_ of MscS was taken to be 7.85*mN/m*^42^. *P*_0.5_ represents the pressure value at which half of the population is in the open state and was determined from the averages of 5-10 traces obtained by using 1-s triangular ramp protocols at the beginning of each experiments (1 *k_B_T /nm*^2^ = 4.114 *mN/m*).

#### Two-State Markov Model

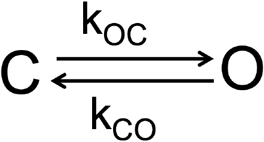

*k_xy_* represents the transition rate, the probability per unit time to make a transition from state Y to state X and is described by the Arrhenius-type relation: 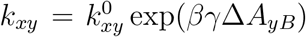 where 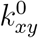 is the intrinsic rate of the system’s attempts to overcome the barrier between states X and Y in the absence of the tension and Δ*A_yB_* is the expansion area from state Y to the barrier, is the applied tension and *β* = 1*/k_B_T*. Equivalently, it can be shown that the transition rates obey detailed balance:

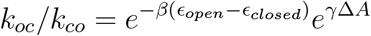

where Δ *A* = Δ *A_cB_* + Δ *A_oB_* is the lateral protein expansion area used in the main text. The detailed balance condition guarantees that once the tension is held fixed the system relaxes to a unique equilibrium distribution where the states are populated according to the Boltzmann distribution. The following parameters were used for the two-state model of MscS: 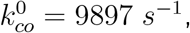, 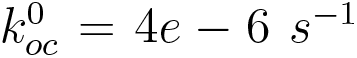, |Δ*A_cB_*| = 7 *nm*^2^ and |Δ*A_oB_*| = 5 *nm*. The tension protocol used in the simulation is the same as the experimental protocol depicted in Fig. 2.

#### Edge Detection

**FIG. 4:**
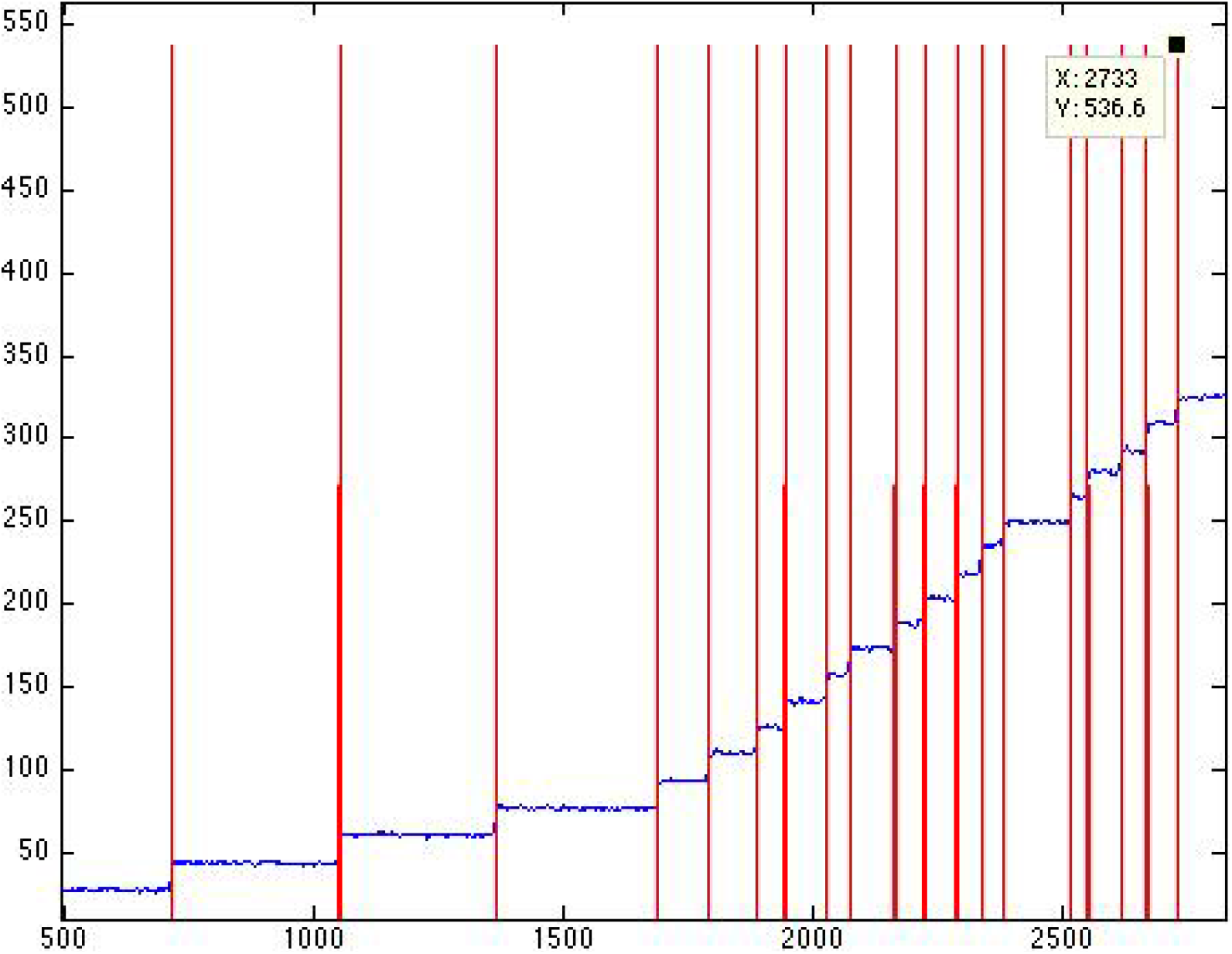
An Edge Detector program (http://cismm.web.unc.edu/resources/tutorials/edge-detector-1d-tutorial/) was employed to detect the single channel events.

## Notes

### Competing Interest Statement

The authors have declared no competing interest.

## References

1 S. Lloyd, Nature 406, 1047 (2000).

2 R. Landauer, IBM journal of research and development 5, 183 (1961).

3 O. J. Maroney, Physical Review E 79, 031105 (2009).

4 L. Gammaitoni, arXiv preprint arXiv:1111.2937 (2011).

5 J. A. Vaccaro and S. M. Barnett, Proceedings of the Royal Society A: Mathematical, Physical and Engineering Sciences 467, 1770 (2011).

6 T. Sagawa, Journal of Statistical Mechanics: Theory and Experiment 2014, P03025 (2014).

7 T. Sagawa and M. Ueda, Phys. Rev. Lett. 102, 250602 (2009).

8 A. B. Boyd and J. P. Crutchfield, Phys. Rev. Lett. 116, 190601 (2016).

9 E. Bormashenko, Entropy 21, 1150 (2019).

10 K. Proesmans, J. Ehrich, and J. Bechhoefer, arXiv preprint arXiv:2006.03240 (2020).

11 K. Proesmans, J. Ehrich, and J. Bechhoefer, arXiv preprint arXiv:2006.03242 (2020).

12 D. H. Wolpert, Journal of Physics A: Mathematical and Theoretical 52, 193001 (2019).

13 C. P. Kempes, D. Wolpert, Z. Cohen, and J. Pérez-Mercader, Philosophical Transactions of the Royal Society A: Mathematical, Physical and Engineering Sciences 375, 20160343 (2017).

14 A. Bérut, A. Arakelyan, A. Petrosyan, S. Ciliberto, R. Dillenschneider, and E. Lutz, Nature 483, 187 (2012).

15 A. Bérut, A. Petrosyan, and S. Ciliberto, Journal of Statistical Mechanics: Theory and Experiment 2015, P06015 (2015).

16 Y. Jun, M. Gavrilov, and J. Bechhoefer, Physical review letters 113, 190601 (2014).

17 M. Gavrilov and J. Bechhoefer, Physical Review Letters 117, 200601 (2016).

18 M. Gavrilov, R. Chétrite, and J. Bechhoefer, Proceedings of the National Academy of Sciences 114, 11097 (2017).

19 J. Hong, B. Lambson, S. Dhuey, and J. Bokor, Science advances 2, e1501492 (2016).

20 L. Martini, M. Pancaldi, M. Madami, P. Vavassori, G. Gubbiotti, S. Tacchi, F. Hartmann, M. Emmerling, S. Höfling, L. Worschech, et al., Nano Energy 19, 108 (2016).

21 R. Gaudenzi, E. Burzurí, S. Maegawa, H. van der Zant, and F. Luis, Nature Physics 14, 565 (2018).

22 O.-P. Saira, M. H. Matheny, R. Katti, W. Fon, G. Wimsatt, J. P. Crutchfield, S. Han, and M. L. Roukes, Physical Review Research 2, 013249 (2020).

23 J. P. Peterson, R. S. Sarthour, A. M. Souza, I. S. Oliveira, J. Goold, K. Modi, D. O. Soares-Pinto, and L. C. Céleri, Proceedings of the Royal Society A: Mathematical, Physical and Engineering Sciences 472, 20150813 (2016).

24 L. Yan, T. Xiong, K. Rehan, F. Zhou, D. Liang, L. Chen, J. Zhang, W. Yang, Z. Ma, and M. Feng, Physical review letters 120, 210601 (2018).

25 P. Moe and P. Blount, Biochemistry 44, 12239 (2005).

26 U. Çetiner, I. Rowe, A. Schams, C. Mayhew, D. Rubin, A. Anishkin, and S. Sukharev, The Journal of general physiology 149, 595 (2017).

27 S. I. Sukharev, W. J. Sigurdson, C. Kung, and F. Sachs, The Journal of general physiology 113, 525 (1999).

28 N. Levina, S. Tötemeyer, N. R. Stokes, P. Louis, M. A. Jones, and I. R. Booth, The EMBO journal 18, 1730 (1999).

29 C. Kung, B. Martinac, and S. Sukharev, Annual review of microbiology 64, 313 (2010).

30 S. Sukharev, B. Martinac, V. Arshavsky, and C. Kung, Biophysical journal 65, 177 (1993).

31 E. Neher and B. Sakmann, Nature 260, 799 (1976).

32 O. P. Hamill, A. Marty, E. Neher, B. Sakmann, and F. Sigworth, Pflügers Archiv 391, 85 (1981).

33 B. Sakmann and E. Neher, Annual review of physiology 46, 455 (1984).

34 B. Martinac, M. Buechner, A. H. Delcour, J. Adler, and C. Kung, Proceedings of the National Academy of Sciences 84, 2297 (1987).

35 An Edge Detector program (http://cismm.web.unc.edu/resources/tutorials/edge-detector-1d-tutorial/) was employed to detect the single channel events.

36 U. Seifert, Reports on progress in physics 75, 126001 (2012).

37 U. Çetiner, O. Raz, S. Sukharev, and C. Jarzynski, Physical Review Letters 124, 228101 (2020).

38 R. Phillips, T. Ursell, P. Wiggins, and P. Sens, Nature 459, 379 (2009).

39 R. Phillips, J. Theriot, J. Kondev, and H. Garcia, Physical biology of the cell (Garland Science, 2012).

40 U. Çetiner, A. Anishkin, and S. Sukharev, European Biophysics Journal 47, 663 (2018).

41 C. Bustamante, J. Liphardt, and F. Ritort, arXiv preprint cond-mat/0511629 (2005).

42 V. Belyy, K. Kamaraju, B. Akitake, A. Anishkin, and S. Sukharev, The Journal of general physiology 135, 641 (2010).

43 Formally, the system has to get back to the same tension value. But this does not make any difference since we can always release the tension instantaneously without changing the work or heat.

44 The software is available at https://qub.mandelics.com.

45 E. R. Kashket, Annual review of microbiology 39, 219 (1985).

46 T. A. Krulwich, G. Sachs, and E. Padan, Nature Reviews Microbiology 9, 330 (2011).

